# Sequence Complexity Dictates Polymer Mixing

**DOI:** 10.64898/2025.12.23.696237

**Authors:** Dibyajyoti Mohanta, Manish Dwivedi, Hiranmay Maity, Debaprasad Giri

**Author notes:** Electronic mail (Dibyajyoti Mohanta).

## Abstract

Polymer association in confined spaces governs diverse phenomena from protein aggregation to DNA condensation. We investigate the sequence-level mechanisms underlying this behavior through exact enumeration of two confined AB-type block copolymers on a two-dimensional lattice, revealing how sequence complexity controls mixing against de-mixing. We reveal that sequence complexity (heterogeneity), quantified by Shannon entropy *H*_2_, acts as a determinant between self-folded low-mixed state and high-mixed state. High-complexity sequences (*H*_2_ = 1.86 bits) with short-length repeats achieve near-complete inter-chain overlap through cooperative chain collapse, while low-complexity blocky sequences (*H*_2_ = 1.26 bits) maintain extended conformations with less overlapping. Free energy analysis reveals a steep increase in mixing barriers for low-complexity compared to surmountable barrier for high-complexity sequences. We find that geometric confinement modulates but does not override these sequence-dependent behaviors, with asymmetric confinement enhancing heterogeneity in overlapping for both sequence type (pronounced in high-*H*_2_ sequences). Our theory could be applicable in assessing the phase behavior of repeat protein and nucleic acid sequences.

## 1. INTRODUCTION

Liquid-Liquid Phase Separation (LLPS) mediated by weak interactions provided the framework for compartmentalization of biomolecules, including protein and RNA, even without any membrane^1^. The formation and properties of such membraneless compartments known as biocondensates depend not only on the overall composition of constituent proteins and nucleic acids but are also sensitive to the underlying sequence patterning of interaction motifs^2,3^. Such sequence-encoded effects exhibit both in single-molecule and multicomponent systems, where sequence composition and patterning shape the conformational landscape of biopolymers^4^. For example, intrinsically disordered proteins (IDPs) and regions (IDRs), which lack well-defined tertiary structures, adopt conformational ensembles directly encoded by their sequences^5,6^. Similarly, nucleic acid sequences also control the formation of secondary structures such as hairpin loops, bulges in RNA, bubbles in DNA, through the specific arrangement of nucleotides^4,7^. Polyampholytes with well-mixed positive and negative charges readily collapse, while polyelectrolytes like poly(A)-RNA or poly(U)-RNA resist condensation under physiological conditions^7,8^. This sequence dependent organization and association of biomolecules serve as a fundamental determinant of phase separation behavior^9^.

In vivo, sequence effects never act in isolation as biomolecules fold and function within crowded, geometrically confined environments that fundamentally alter their behavior^10,11^. The cellular environment contains 20 − 40% macromolecules by volume, creating steric constraints that restrict conformational entropy and modulate interaction energetics^12,13^. Recent studies have demonstrated that confinement can reduce critical concentrations for phase separation by 4-10 fold, making LLPS thermodynamically accessible at physiological protein concentrations^14,15^. Confinement creates a delicate balance: it restricts the conformational entropy of RNA and proteins, while at the same time enhancing local concentration effects that strengthen intermolecular interactions^16,17^. Understanding how geometric confinement couples with sequence patterning to determine phase separation driven multi-polymer association is essential.

Biomolecule aggregation often features hierarchy of assembly that begins with monomers, dimers, oligomers, which subsequently grow into network-like topologies characteristic of condensates beyond the critical density^4,18–^. In this context, we leverage insights from previous theoretical and simulation studies on confined two-polymer systems (dimers) to identify the key factors that control their folding, mixing, and segregation^21–28^. However, the majority of these studies have focused on homopolymers, which are largely dominated by entropic effects^24,26^. By design, these models neglect the sequence-specific enthalpic interactions that drive the microphase separation essential in facilitating polymer association^29^. Thus, a fundamental gap remains in our understanding of how sequence patterning and geometric confinement jointly regulate the initial association of polymers. To remedy this, we present an exactly solvable minimal model that captures interplay between sequence complexity, geometric confinement, and polymer association, through exhaustive enumeration.

A central challenge in predicting sequence-dependent polymer interactions is the quantitative characterization of sequence patterning. Shannon entropy, calcu-lated from the probability distribution of n-letter words (n-grams), provides a robust measure of sequence patterning defined as sequence complexity. It correlates strongly with conformational heterogeneity and phase separation propensity^19,30–32^. To this end, we employ a system of two AB-type block copolymers where sequence complexity is quantified by the bi-gram entropy, *H*_2_. This approach is motivated by recent studies on block copolymer systems composed of two distinct bead types, investigating sequence-controlled self-assembly and aggregation^29,33^.

This study unfolds in three key parts. First, we map out the role of sequence complexity and confinement anisotropy in shaping mixing of two polymers. Next, we in-depth analyze the free energy landscape of mixing for different *H*_2_ sequences within varying confinement. Finally, we explore how sequence complexity affects the actual size of two-polymer systems during the overlapping pathway. We conclude our study by investigating the mixing behavior for a diverse pair of sequences.

## II. MODEL AND METHOD

We model a system of two block co-polymers (BCPs), each consisting of *N* + 1 = 18 beads, as *self(mutual)-attracting-self(mutual)-avoiding walks (S(M)AS(M)AW)* within a symmetric confinement *S*_*s*_ with side length *L*_*xs*_ = *L*_*ys*_ = 10 lattice units, and an asymmetric con-finement *S*_*r*_ with *L*_*xr*_ = 20 and *L*_*yr*_ = 5 lattice units. Both confinements have an equal area of 100 square lattice units, maintaining a constant volume fraction of (2 × 18)*/*100 = 0.36 [Fig. 1]. We define a specific, attractive interaction *ϵ*_*sp*_ = *ϵ* between non-bonded nearest-neighbor beads of same type (*A* − *A* or *B* − *B*). Interactions between different beads type (*A* − *B*) termed as non-specific contacts and are set to zero (shown as × in Fig. 1).

**FIG. 1.**
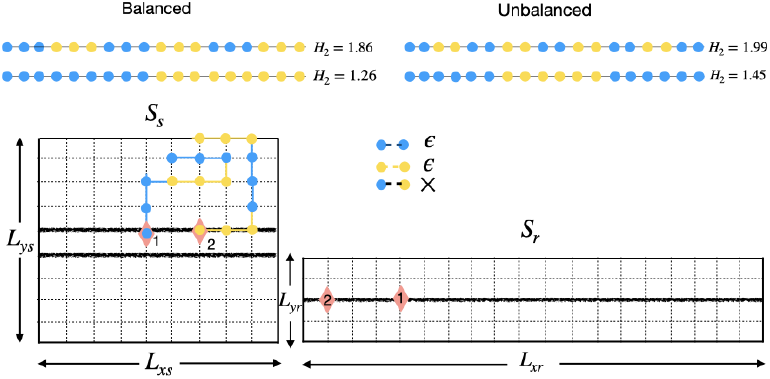
Schematic diagram of the two block co-polymer (BCP) model. (Top) Examples of balanced and unbalanced BCP sequences with A-type marked as blue and B-type as yellow. The sequence complexity (*H*_2_) is shown for each pair. (Bottom) The two BCPs are confined within following geometries: a square box (*S*_*s*_) of 10× 10 and a rectangular box (*S*_*r*_) of 20 × 5 lattice units. The shaded middle layer indicates the set of possible starting positions for the polymers. In *S*_*s*_, chains are enumerated from one of the two shaded layers, as 5^*th*^ and 6^*th*^ layers (*L*_*ys*_) are mirror images to each other. Pink diamonds represent an example “origin-pair” for the two polymers (labeled 1 and 2). Strength of interaction between same type of beads is taken as *ϵ*, whereas non-specific interaction strength is zero.

### A. Sequence Entropy *H*_2_ of Block copolymers

The n-gram entropy measures the randomness of overlapping subsequences (‘words’) in a polymer chain, where beads are treated as letters (e.g., A/B for a diblock copolymer or hydrophobic/polar in HP models). Following Herzel^34^, the bi-gram (2-letter word) entropy *H*_2_ for a given 18-letter sequence of A and B is computed from the frequencies of all overlapping 2-letter words, yielding 17 such bi-grams per sequence.

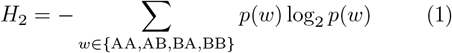

where the probability of each bi-gram *w* in the 18-bead sequence is estimated as *p*(*w*) = *n*(*w*)*/*17, with *n*(*w*) being the count of that bi-gram. All four bi-grams are equal-a-priori, but their empirical probabilities *p*(*w*) reflect the actual bi-gram composition of the specific sequence^35^.

Our study focuses on four distinct two-BCP systems characterized by their sequence complexity *H*_2_ and over-all A/B composition (Table I). Each system consists of a sequence listed in Table I together with its complementary partner obtained by exchanging A and B at every position. For example, the system *H*_2_ = 1.86 consists of the AAABBBAAABBBAAABBB sequence and its complementary partner BBBAAABBBAAABBBAAA with *A/B* = 1 in individual sequence and overall two-BCP system.

**TABLE I.**
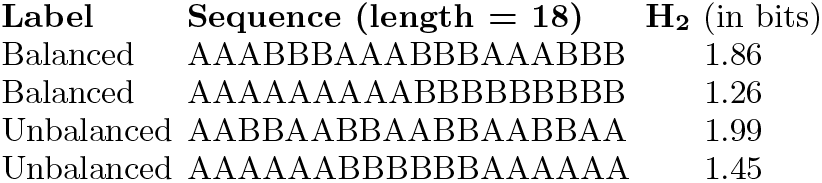
Bi-gram entropy *H*_2_ (in bits) for four designed 18-bead polymer sequences composed of A and B beads. “Balanced” sequences have equal numbers of A and B beads, whereas “Unbalanced” sequences have unequal A/B composition. Larger *H*_2_ indicates greater sequence complexity.

In balanced systems (*H*_2_ = 1.86 and *H*_2_ = 1.26), each chain has equal numbers of A and B beads, while in unbalanced systems (*H*_2_ = 1.99 and *H*_2_ = 1.45), the chains have unequal A/B composition. Although many distinct sequences share similar *H*_2_ values, this work focuses on four representative systems chosen to represent the high and low ends of the complexity spectrum, from short periodic patterns to more blocky ones, with *H*_2_ values ranging approximately from 1.99 to 1.26 bits.

### B. Enumeration procedure

The exact partition function of this binary BCP system can be calculated by exact enumeration of all conformations of two polymer chains of length *N* = 17 (i.e., *N* + 1 beads)^36^. At this fixed chain length and volume fraction (*ϕ*_*c*_ = 0.36), the full conformational space is already large, so extending to longer chains would be computationally expensive^26,36^.

To obtain reliable statistics at fixed *N* without increasing chain length, we systematically scan the accessible volume by varying the positions of the polymer origins inside the confinement. Specifically, we restrict the starting positions (origins) of both polymers to the middle layer of the box, as indicated by the shaded line in [Fig. 1]. For each possible unique pair of origin positions on this line (an “origin-pair”), we enumerate all conformations of the two BCPs; providing an exhaustive conformational ensemble for each origin-pair.

This procedure yields 10 × 9 = 90 distinct origin-pairs for the *S*_*s*_ box and 20 × 19 = 380 origin-pairs for the *S*_*r*_ box. Throughout the study, observables are reported both as averages over individual origin-pairs and as ensemble averages over all origin-pairs, thereby ensuring a uniform scan of the confinement’s effect while remaining computationally feasible.

The system is evaluated in the canonical ensemble. For a origin-pair, the partition function *Z* is computed as a sum over all possible conformations:

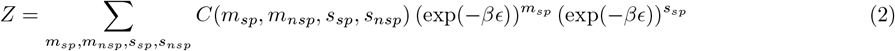

Here, *C*(*m*_sp_, *m*_nsp_, *s*_sp_, *s*_nsp_) denotes the number of conformations with:

*m*_**sp**_: inter-polymer specific contacts (A–A or B–B between polymers)

*m*_**nsp**_: inter-polymer non-specific contacts (A–B between polymers)

*s*_**sp**_: intra-polymer specific contacts (A–A or B–B within each polymer, summed)

*s*_**nsp**_: intra-polymer non-specific contacts (A–B within each polymer, summed)

In this model (see [Fig. 1]), only specific contacts are energetically weighted (i.e. *ϵ*_*nsp*_ = 0). So, the system energy depends solely on the total number of specific contacts (*m*_sp_ + *s*_sp_) (Eq. 2). On the other hand, polymer overlap or mixing involves contributions from both specific (*m*_*sp*_) and non-specific (*m*_*nsp*_) contacts. *β* = 1*/*(*k*_*B*_*T*) is the inverse temperature. For this study, we used reduced units where the Boltzmann constant *k*_*B*_ is set to 1 and the temperature T is fixed at 0.5. To calculate the overall ensemble average for a given sequence type, we followed Eq. 2 across all origin pairs.

## III. RESULTS

### 1. Statistical Mechanics of Contact Distribution

Multiple polymer assemblies such as condensates formed by LLPS, chromatin organization, and crowded macromolecular complexes, exhibit network-like architectures that typically rely on strength of multivalent inter-chain interactions^4,37^. In [Fig.2], we analyze how sequence complexity and confinement jointly control BCP mixing by examining the fraction of inter-chain contacts per polymer, *ϕ*_*nc*_ = *N*_*c*_*/*(*N* +1) with *N*_*c*_ = *m*_sp_+*m*_nsp_, for high-complexity (*H*_2_ = 1.86, short-length repeats) and low-complexity (*H*_2_ = 1.26, blocky) balanced sequences under symmetric (*S*_*s*_, panels a,b,c) and asymmetric (*S*_*r*_, panels d,e,f) confinement. Each radially outward bar in the polar plots (Fig.2) represents *ϕ*_*nc*_ for a distinct origin pair of the two BCPs within the confinement.

At weak interaction strength *ϵ* = −0.5 [Fig.2 (a), (d)], both sequences exhibit a modest and comparable overlap (*ϕ*_*nc*_ < 0.5) with *H*_2_ = 1.26 being slightly higher. This indicates that conformational entropy dominates over low specific interaction energy which is insufficient to elicit sequence-dependent differences^38^. As attraction strength increases to *ϵ* = −1.0, the distributions bifurcate [Fig. 2(b,e)]. Polymers with higher *H*_2_ display a much broader spread of *ϕ*_*nc*_ values and access configurations with substantially higher overlap. Under strong attractive condition *ϵ* = −3.0 [Fig. 2(c,f)], the distinction is maximized. *H*_2_ = 1.86 sequences approach near-complete overlap (*ϕ*_*nc*_ ∼ 1.1–1.2), whereas blocky *H*_2_ = 1.26 sequences saturate at significantly lower *ϕ*_*nc*_ values (≈ 0.6), in both confinements^30,33^.

**FIG. 2.**
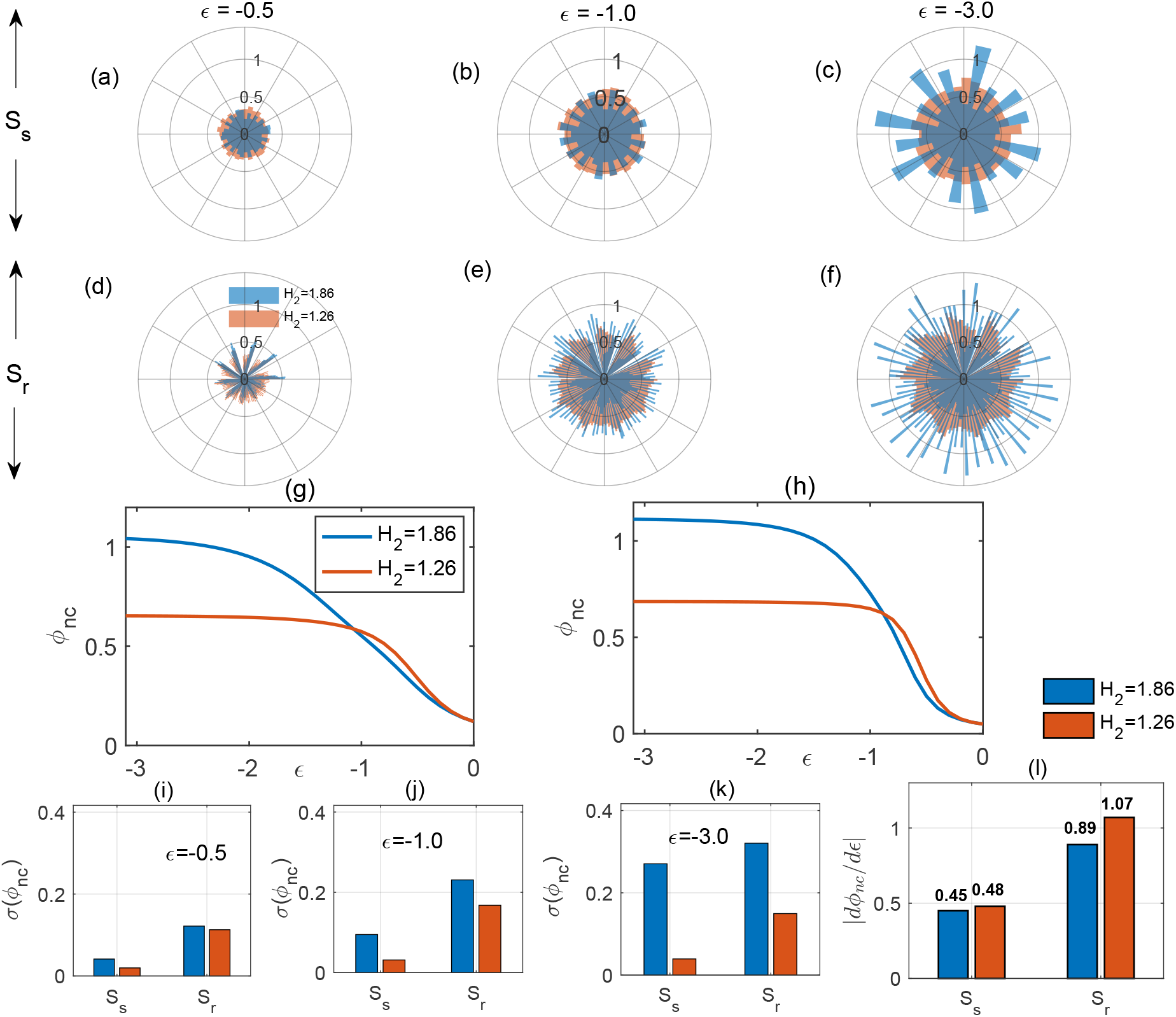
Sequence-dependent inter-chain overlapping across origin pairs for balanced sequences. Polar plots of fraction of overlap contacts (*ϕ*_*nc*_ = *N*_*c*_*/*(*N* + 1)) for all origin-pairs at interaction strengths *ϵ* = −0.5, −1.0, −3.0 (a,b,c) for high-complexity (*H*_2_ = 1.86), and (d,e,f) for low-complexity (*H*_2_ = 1.26) sequences in square confinement *S*_*s*_ and rectangular confinement *S*_*r*_. Compared to *S*_*s*_, *S*_*r*_ shows more heterogeneity in *ϕ*_*nc*_, especially in *S*_*r*_ and at stronger *ϵ*. Ensemble-averaged ⟨*ϕ*_*nc*_⟩ over origin pairs versus interaction strength *ϵ* in *S*_*s*_ (g) and *S*_*r*_ (h), showing higher mixing for high-*H*_2_ at strong interactions for both confinements. Heterogeneity in mixing, quantified as standard deviation of *ϕ*_*nc*_ across origin-pairs in *S*_*s*_ and *S*_*r*_ (i, j,k for different values of *ϵ*), highlighting enhanced variability in high-*H*_2_ and *S*_*r*_. Initial mixing rates *dϕ*_*nc*_*/d* |*ϵ*| in the linear regime (l), revealing faster but shallower response for low-*H*_2_, while slow but progressive mixing for high–*H*_2_ (pronounced in *S*_*r*_).

Ensemble-averaged *ϕ*_*nc*_ over all origin-pairs, reinforce these trends [Fig. 2(g,h)]. At low |*ϵ*|, *H*_2_ = 1.26 exhibit mild higher *ϕ*_*nc*_ than *H*_2_ = 1.86 sequences, reflecting their greater sensitivity to weak attractions. However, as |*ϵ*| increases, high-*H*_2_ sequences overtake low-*H*_2_ sequences in both confinements, achieving much higher saturation overlap. The contiguous arrangement of identical beads in low *H*_2_ blocky chains make intra-chain folding entropically favorable, allowing the BCPs to satisfy most of their valency via compact self-collapsed states (see [Fig. S1] in SI) ^39–41^. In contrast, the high complexity *H*_2_ sequences make intra-chain folding entropically costly due to short-repeat lengths, leaving attractive beads exposed to form the multivalent inter-chain mixing^19,30,42^.

To quantify how confinement anisotropy modulates the heterogeneity in fraction of inter-contact distribution, we computed the standard deviation of *ϕ*_*nc*_ across all origin-pairs at different strengths [Fig. 2(i,j,k)]. At all interaction strengths, both type of sequences within *S*_*r*_ exhibits higher heterogeneity than *S*_*s*_, reflecting the broader spectrum of accessible conformations and overlap states in anisotropic confinement^22,43^. Moreover, among the sequences, high-complexity sequences consis-tently show larger heterogeneity than their low-*H*_2_ counterparts in both geometries, indicating richer landscape of partially and highly mixed configurations for low and high complexity sequences respectively.

The linear fits to *ϕ*_*nc*_ in the linear regime (low |*ϵ*|) reveal distinct “mixing rates” for the two sequence types [Fig. 2(l)]. Low-*H*_2_ sequences display a steeper initial slope, presenting a faster but shallow response to increasing attraction that quickly saturates at low overlap. In contrast, high-*H*_2_ sequences exhibit a more moderate initial increase but ultimately progress to complete overlap, with effective mixing rates |d*ϕ*_*nc*_*/*d*ϵ*| ≈ 0.45 in *S*_*s*_ and ≈ 0.89 in *S*_*r*_. Thus, asymmetric confinement nearly doubles the mixing rate for high-complexity sequences by pre-aligning chains and reducing the entropic penalty of association^44^. This is consistent with prior studies showing that confinement can shift phase boundaries and reduce the critical concentration required for LLPS ^14,15,44,45^.

To explore the molecular origins of sequence complexity-dependent overlapping behavior, we present the fraction of total specific contacts *ϕ*_*sp*_ = (*m*_*sp*_ + *s*_*sp*_)*/*(*N* + 1) in [Fig. 3 (a)]. A key finding is that low-*H*_2_ = 1.26 sequences consistently form a higher fraction of *ϕ*_*sp*_ compared to their high-*H*_2_ counterparts across both confinements. This observation appears counterintuitive at first, as one might expect that a greater number of specific contacts would promote enhanced mixing for *H*_2_ = 1.26. However, this can be explained by decomposing total specific contacts into their inter- and intra-chain components expressed as *ϕ*_*msp*_ = *m*_*sp*_*/*(*N* +1) and *ϕ*_*ssp*_ = *s*_*sp*_*/*(*N* + 1) in [Fig. 3 (b) and (c)] respectively. Blocky, low-*H*_2_ sequences sequences have contiguous blocks of identical beads (more homotypic contacts per segment), making intra-chain folding energetically favorable (with higher *ϕ*_*ssp*_ values for *H*_2_ = 1.26 in [Fig. 3 (c)]. This reduces inter-chain overlap driven by specific contacts defined by *ϕ*_*msp*_ [Fig. 3 (b)] for *H*_2_ = 1.26. In sharp contrast, short-length repeats in high-*H*_2_ chains maintain more open conformations because intra-chain folding incurs a large entropic penalty when attempting to bring distant non-contiguous attractive beads into contact ^19,33^. This leaves many attractive sites exposed and available to form inter-chain-specific contacts featuring higher *ϕ*_*msp*_ for *H*_2_ = 1.86 in [Fig. 3 (b)]. This highlights how low-*H*_2_ sequences are energetically favored (via more total contacts) but entropically disfavored (due to self-folding) for inter-chain mixing, whereas high-*H*_2_ sequences trade some energetic gains for entropic advantages through enhanced inter-chain association [Fig. 3 (b)]^41^. These trends align with studies demonstrating that strong homotypic interactions in proteins promote self-association over intermolecular mixing^42^.

**FIG. 3.**
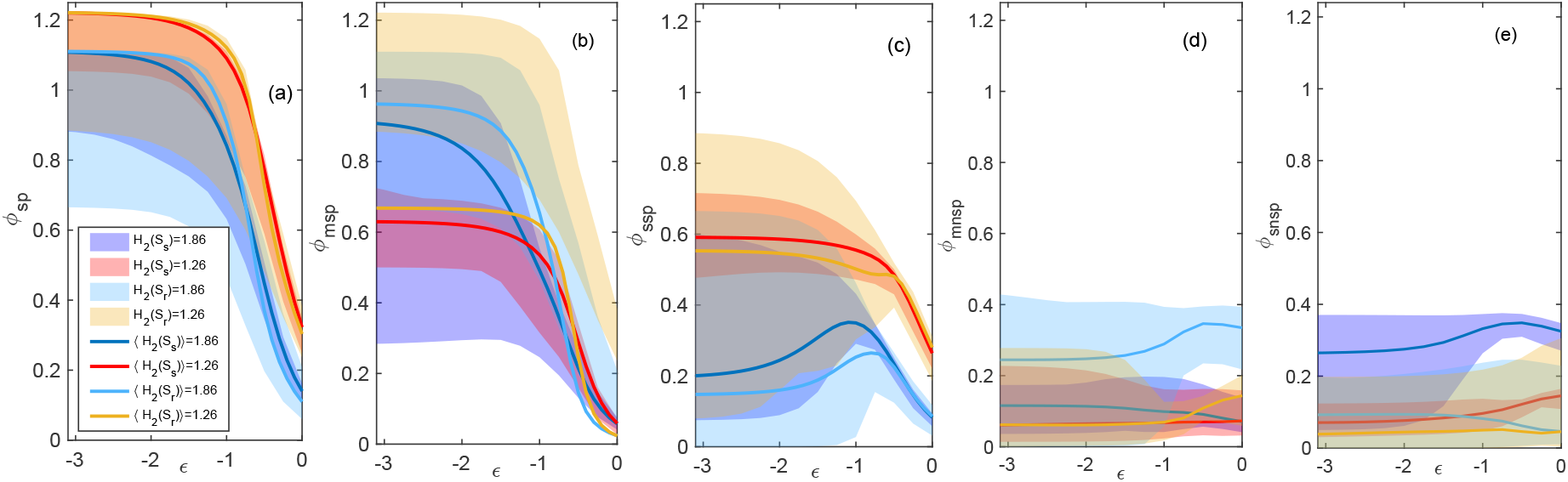
Decomposition of contact fractions reveals sequence-dependent preferences for self-association versus mixing. (a) For both confinements total phase-separated (i.e. *ϕ*_*sp*_ = *ϕ*_*msp*_ + *ϕ*_*ssp*_) contact fraction are higher for *H*_2_ = 1.26 than *H*_2_ = 1.86, representing *H*_2_ = 1.26 energetically dominated. (b) Inter-chain specific contact fraction *ϕ*_*msp*_ are higher for high-*H*_2_, promoting sequence-specific mixing. (c) Intra-chain specific contact fraction *ϕ*_*ssp*_ dominate for low-*H*_2_, favoring self-association. (d) Inter non-specific contact fraction *ϕ*_*mnsp*_ (A-B) are higher for high-*H*_2_, contributing to overall overlap. (e) Intra-chain non-specific contact fraction *ϕ*_*snsp*_ remain minimal for both sequences.

The fraction of inter-chain non-specific contacts (*ϕ*_*mnsp*_ = *m*_*nsp*_*/*(*N* + 1)), which do not contribute to the Boltzmann weight but still takes part in mixing, is low overall but also remains higher for short-range sequences (*H*_2_ = 1.86) in both confinements [Fig. 3 (d)]. This arises because, in high-*H*_2_ chains, non-interacting blocks interspersed between favorable interaction blocks can remain overlapped between two chains due to the reduced entropic cost of constraining these short segments ^40^. This behavior mirrors observations in sticker-spacer models of LLPS, where short spacers tend to remain in proximity due to the strong attachments of adjacent stickers ^19,40^. Finally, intra-chain non-specific contacts (*ϕ*_*snsp*_ = *s*_*nsp*_*/*(*N* + 1)) provide minimal contributions to folding, thereby emphasizing the dominance of energy-driven specific interactions [Fig. 3(e)]. Collectively, Figs. 2 and 3 demonstrate that the spatial arrangement of interaction sites, rather than their total number or binding strength, governs whether polymers assemble into self-collapsed or form cooperative polymer organization^30,33^.

### 2. Free Energy Profile Analysis

To determine the thermodynamic driving forces governing sequence-dependent mixing, we analyzed the free energy profiles *F* (*N*_*c*_), calculated from partition function with *N*_*c*_ inter-chain contacts *Z*(*N*_*c*_) [Figure 4]:

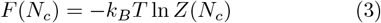

where *Z*(*N*_*c*_) is expressed as ^22^,

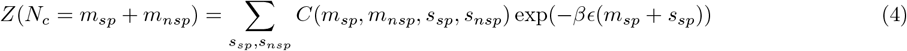

**FIG. 4.**
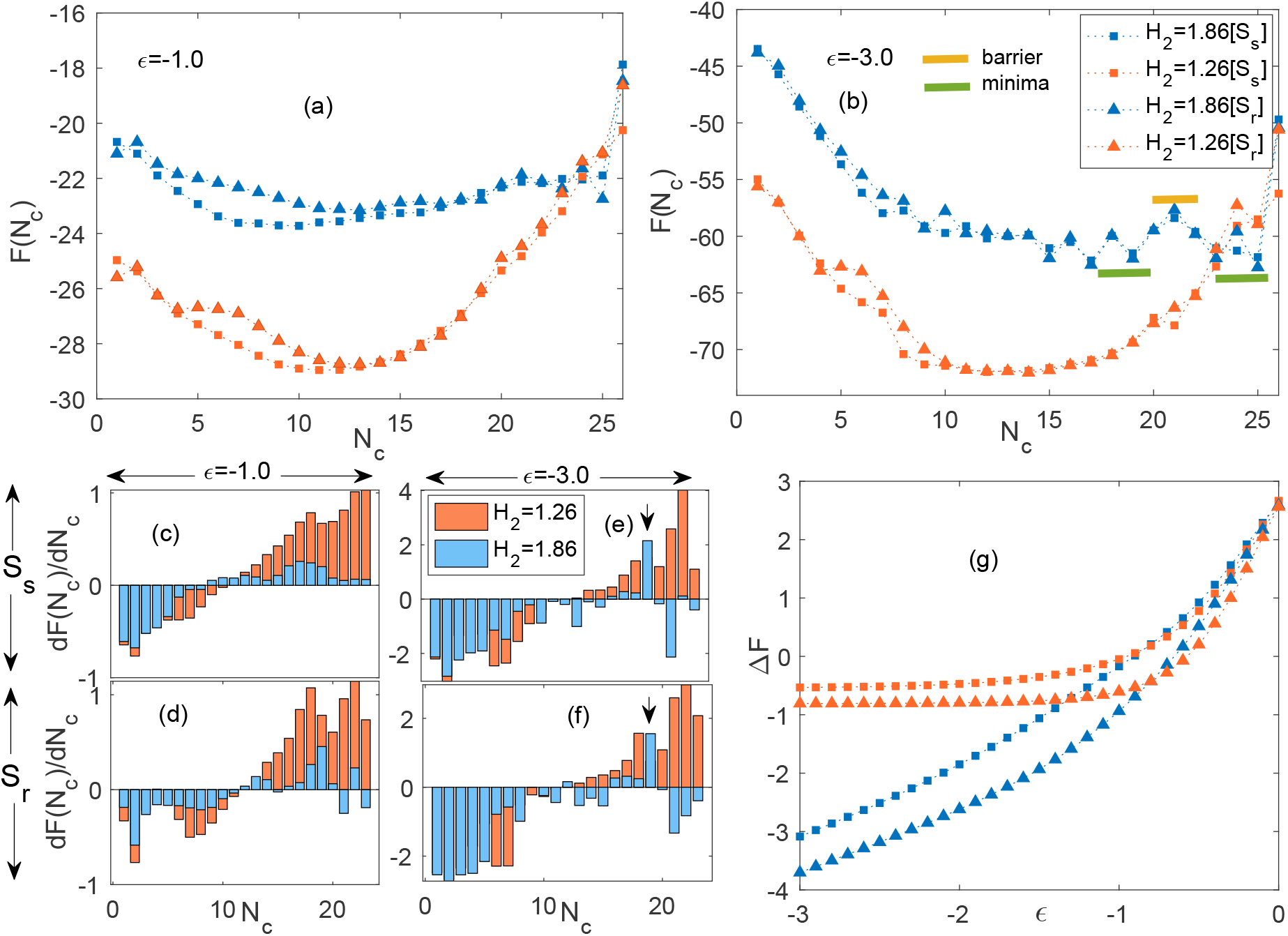
Free energy landscapes and barriers reveal the thermodynamic basis for sequence-dependent mixing: (a, b) Free energy profiles *F* (*N*_*c*_) as a function of inter-chain contacts *N*_*c*_ at weak (*ϵ* = −1.0) and strong (*ϵ* = −3.0) interaction strengths, respectively, for high-complexity (*H*_2_ = 1.86, blue) and low-complexity (*H*_2_ = 1.26, red) sequences in square (*S*_*s*_) and rectangular (*S*_*r*_,triangles) confinements. High-*H*_2_ sequences exhibit an initial entropic penalty at low *N*_*c*_ but a favorable crossover to lower free energy at high *N*_*c*_, indicative of a surmountable barrier leading to stable mixed states. The barrier is marked as yellow bar separating the two minimas marked as green bars in (b). Whereas, low-*H*_2_ sequences show a monotonic increase, disfavoring mixing. (c,d,e,f) Local gradients *∂F/∂N*_*c*_ (in units of *k*_*B*_ *T*) highlighting barriers and driving forces: (c, e) at *ϵ* = −1.0 for *S*_*s*_ and *S*_*r*_; (d, f) at *ϵ* = −3.0 for *S*_*s*_ and *S*_*r*_. High-*H*_2_ sequences display a characteristic spike around *N*_*c*_ ≈ 20 (downward arrows), marking the barrier between demixed and mixed minima, with negative gradients post-barrier favoring mixing more pronounced in asymmetric *S*_*r*_ confinement. Low-*H*_2_ sequences show persistently positive gradients, resisting overlap. (g) Free energy difference Δ*F*_mixing_ between mixed and demixed states as a function of interaction strength *ϵ*, confirming cooperative transitions to mixing for high-*H*_2_ (decreasing Δ*F*) versus persistent demixing for low-*H*_2_ (positive Δ*F*) in both confinements

At both weak [Fig. 4 (a)] and strong [Fig. 4 (b)] interaction strengths, high-*H*_2_ exhibit higher free energy in the low to moderate contact regime (*N*_*c*_ < 20) compared to low-complexity sequences. This initial penalty arises because the *H*_2_ = 1.86 chains must overcome a translational and orientational entropic penalty to align due to their short repeats, which is not the case for low-*H*_2_ blocky chains^46,47^. However, when sufficient mixing (*N*_*c*_ > 20) has taken place in high-*H*_2_ sequences, the network of inter-chain contacts becomes sufficiently connected (enthalpy dominated) to overcome the entropic cost of chain alignment. So, the threshold which arises near *N*_*c*_ ≈ 20 indicates the surmountable free energy barrier, beyond which the free energy landscape inverts 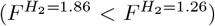, indicating that the highly mixed network becomes the thermodynamic ground state (global minima) for high-*H*_2_^48^. The free energy barrier (yellow bar) and the low-mixed and high-mixed (global) minima marked as green bars are shown in [Fig. 4 (b)]. This mixing threshold at (*N*_*c*_ ≈ 20) basically corresponds to the valley region in density of state *D*(*N*_*c*_) between the double-peaks [Fig.S2 (c) and (g) in SI]. In contrast, low-*H*_2_ sequences cross their free energy minima at *N*_*c*_ ≈ 15 and beyond higher mixing incurs elevated free energy (unfavorable), indicating their unimodal nature in *D*(*N*_*c*_) [Fig. S2 (b),(f) (d) and (h) in SI].

To quantify the resistance to mixing, we computed the local gradient of the free energy landscape, *∂F/∂N*_*c*_ at *N*_*c*_ [Fig. 4 (c)-(f)]. A steep positive gradient indicates a strong thermodynamic penalty against mixing, while a flat or negative gradient implies accessible transitions^49^. In both confinements, the local gradient is negative at low *N*_*c*_ due to the fact that the confinement itself gives stability to low mixed states^44^. There is a crossover of *∂F/∂N*_*c*_ from negative to positive values at *N*_*c*_ ≈ 12 – 13 indicating beyond this point overlapping or mixing requires overcoming an increasing thermodynamic penalty. At low strength *ϵ* = −1.0, *H*_2_ = 1.26 sequence’s *∂F/∂N*_*c*_ lead over high-*H*_2_ sequences in both *S*_*s*_ and *S*_*r*_ at high *N*_*c*_ regime indicating their high resistance in mixing. At high strength *ϵ* = −3.0, [Fig. 4 (e),(f)], after crossing zero in both *S*_*s*_ and *S*_*r*_, high *H*_2_ = 1.86 sequences constantly feature low gradient values until a spike in *∂F/∂N*_*c*_ ≈ 20 (marked by vertical down arrow), indicating the free energy barrier or mixing threshold separating the minimas in [Fig. 4 (b)]. Beyond this peak local gradient, we see *∂F/∂N*_*c*_ going near-zero or negative again (especially in *S*_*r*_) showing the propensity of mixing of high-*H*_2_ sequences. The effect of confinement is also apparent since the local peak gradient in *S*_*s*_ is 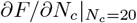 is 2.15 *k*_*B*_*T* while for *S*_*r*_, it is 1.55 *k*_*B*_*T* making *S*_*r*_ the preferred environment for mixing. While low-*H*_2_ sequences show maximum local gradient at higher *N*_*c*_ revealing the mixing is highly unfavorable. The negative *∂F/∂N*_*c*_ values on both sides of the spike for high-*H*_2_ sequences indicate a double-well free energy landscape. In such landscapes, the system can undergo spontaneous transitions between basins (mixed to demixed) via thermal fluctuations ^50,51^. Song et al.^52^ reported such results as tau-RNA complexes reversibly cross LLPS boundaries, with sequence charge patterns tuning the effective barrier between dilute and dense phases.

We further quantified the thermodynamic preference for the mixed state by calculating the global free energy difference Δ*F*_mixing_ between the mixed 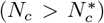 and less-mixed phases [Fig. 4 (g)]. The probability of the mixed phase is given by the normalized partition sum:

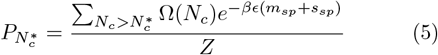

where 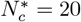 is the identified nucleation threshold [Fig. S2, SI] and used for both low and high *H*_2_ sequences. The relative stability is then:

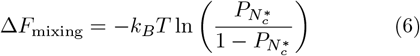

Δ*F*_mixing_ for *H*_2_ = 1.26 remains positive (disfavoring mixing) even at high |*ϵ*| before saturating just below 0. However, the Δ*F*_mixing_ value for the *H*_2_ = 1.86 sequences drops gradually and reaching ∼ −3 for *S*_*s*_ and ∼ −4 for *S*_*r*_ at *ϵ* = −3, confirming a cooperative transition to the mixed phase ^43^.

### 3. Effect of sequence complexity on polymer size

The joint probability distribution *P* (*R*_end_, *N*_*c*_), over all origin pairs with *R*_end_ as the end-to-end distance of polymer 1 within the binary polymer system, reveals sequence-dependent coupling between chain compactness and intermolecular contacts [Fig. 5]. At weak interactions (*ϵ* = −1.0), both sequences in *S*_*s*_ exhibit low-probability distributions (*p* ≈ 0.05 − 0.1) spanning moderate *R*_end_ (4 − 10) and *N*_*c*_ (10 − 22), indicating limited mixing [Fig. 5 a,b]. In *S*_*r*_, high-*H*_2_ show emerging heterogeneity with broader *R*_end_ (10 − 20), consistent with anisotropy enhancing alignment opportunities as seen in contact heterogeneity [Fig. 2] and broader *R*_end_ distributions (*SI* [Fig. S1 e])^43,45^.

**FIG. 5.**
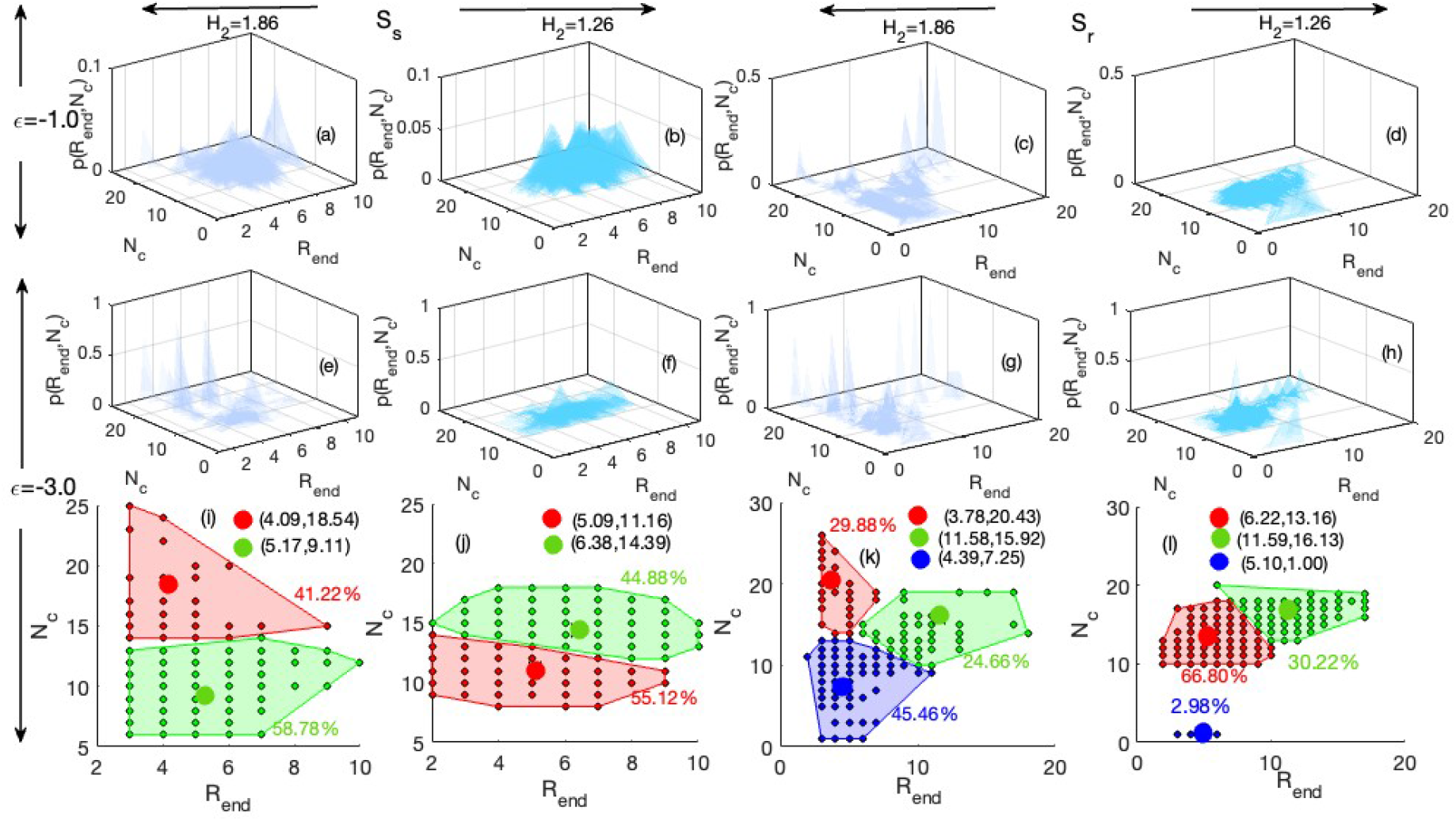
Joint probability distributions *P* (*R*_**end**_, *N*_*c*_) reveal sequence and confinement dependent conformational ensembles. Top: Distributions at *ϵ* = −1.0 (a-d) and Middle: at *ϵ* = −3.0 (e-h) with *S*_*s*_ (square; a,b,e,f) and *S*_*r*_ (rectangular; c,d,g,h). Bottom: k-means clusters at *ϵ* = −3.0 (i-l), with centroids (circles) and colors denote clusters. High-*H*_2_ shows discrete basins favoring low *R*_end_/high *N*_*c*_, indicative of co-collapse; low-*H*_2_ remains extended compared to *H*_2_ = 1.86 with moderate *N*_*c*_.

At strong interactions (*ϵ* = −3.0), high-*H*_2_ sequences display discrete conformational clusters, echoing double-well free energy landscapes [Fig. 4 (b)] that enable cooperative transitions to mixed states. At higher strength, *H*_2_ = 1.86 [Fig. 5 (e),(g)] promotes mixing as well as collapse with higher *P* (*R*_end_, *N*_*c*_) than *H*_2_ = 1.26 [Fig. 5 (f),(h)]. For better identification of mixed-collapsed phase population, we analyze *P* (*R*_end_, *N*_*c*_) by k-cluster analysis, where we focus on high mixed cluster. In *S*_*s*_ [Fig. 5 (i)], k-means clustering (k=2) identifies a highly mixed cluster (41.22%) at low *R*_end_ (4.09 *±* 0.8), indicating inter-chain-driven co-collapse. In *S*_*r*_ [Fig. 5 g,k], k=3 reveals greater heterogeneity, with the *H*_2_ = 1.86 sequences exhibit high-mixed (*N*_*c*_ = 21.43 *±* 1.78) cluster centroid at *R*_end_ = 3.78 *±* 0.4, confirming anisotropy amplifies sequence-encoded mixing without overriding it. However, low-*H*_2_ sequences (*H*_2_ = 1.26) remain largely unimodal in *S*_*s*_ [Fig. 5 f,j]; centered at moderate *R*_end_ (7 − 10) and *N*_*c*_ (10 − 15), resisting compaction due to self-association. In *S*_*r*_ [Fig. 5 h,l] k=3, confinement induces mild heterogeneity, but higher *N*_*c*_ cluster occurs at extended *R*_end_ (11.59 *±* 1.61, *N*_*c*_ = 16.13 *±* 1.00), suggesting expansion to accommodate contacts which is opposite to high-*H*_2_ co-collapse^33,40^. This shows the mixing pathway of high-*H*_2_ sequences exhibit co-collapse state while low-*H*_2_ remains in low-mix-expanded state.

### 4. Unbalanced Sequence Analysis

The extension to unbalanced sequences *H*_2_ = 1.99 and *H*_2_ = 1.45 [Fig. 1], where the ratio of beads type in each polymer chain deviates from unity (*A/B* 1), confirms the robustness of complexity-dependent behavior across distinct composition (stoichiometry)^29^. We compared ensemble average of fraction of inter-chain contact formation /mixing between a high-complexity unbalanced sequence (*H*_2_ = 1.99) with a low-complexity unbalanced sequence (*H*_2_ = 1.45) illustrated in [Fig 6 (a) and (b)].

**FIG. 6.**
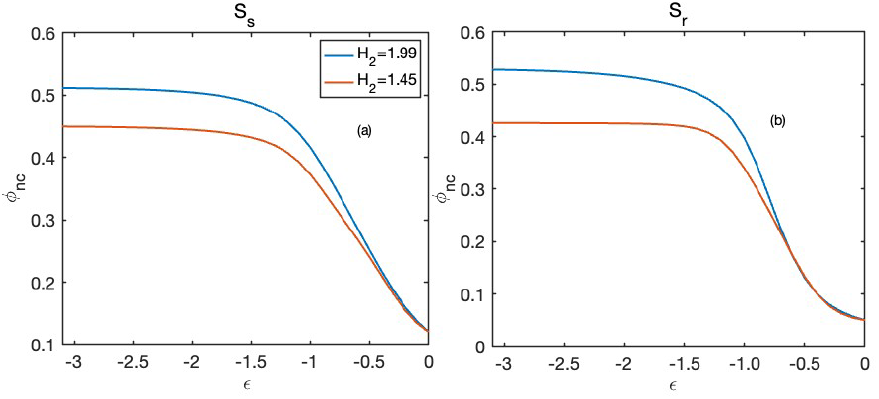
Sequence complexity governs mixing across compositions. Unbalanced sequences with higher *H*_2_ = 1.99 ( blue) exhibit increased inter-chain overlapping compared to low-*H*_2_ = 1.45 (orange) in both square (*S*_*s*_) and rectangular (*S*_*r*_) confinements. However, the absolute fraction of interchain contacts is systematically reduced for unbalanced sequences relative to balanced high-*H*_2_ sequences.

**FIG. 7.**
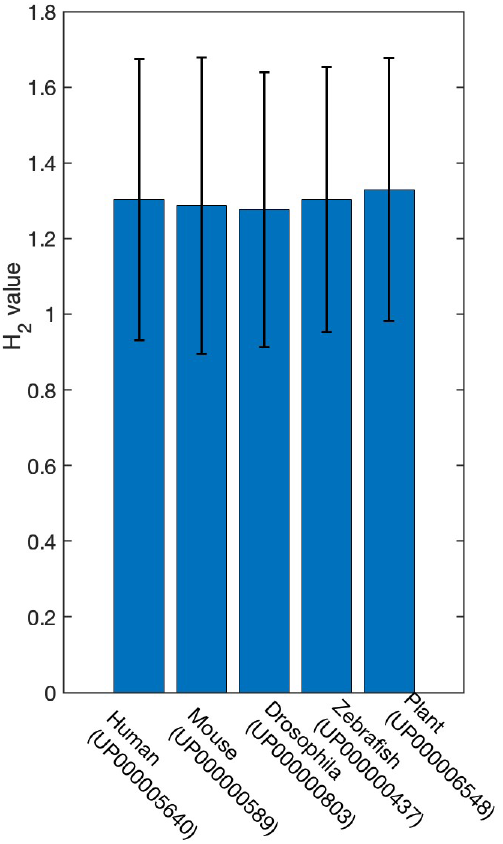
Shannon bi-gram entropy (*H*_2_) of poly-XY regions across eukaryotic proteomes. (Human, Mouse, *Drosophila, Zebrafish*, and Plant). All proteomes exhibit clustered *H*_2_ values (mean ≈ 1.29 bits) with comparable interquartile ranges. Natural polyXY sequences fall between our designed low-complexity (*H*_2_ = 1.26) and high-complexity (*H*_2_ = 1.86) variants, occupying an intermediate mixing regime.

Importantly, while overall mixing levels are reduced for unbalanced compared to balanced sequences, still high-*H*_2_ = 1.99 continues to favor mixing 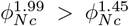 in both confinements. This reduction arises because unbalanced sequences have a higher translational entropy than balanced ones, thereby resulting less alignment driven mixing^53^. Our observation that ‘unbalanced’ stoichiometries (*A* ≠ *B*) reduce polymer mixing warrants careful distinction from the ‘magic-ratio effect’ recently established for multicomponent condensates^53^. In systems driven by heterotypic interactions (e.g., sticker-spacer models where A binds B), phase separation is often suppressed at 1:1 stoichiometry because the formation of small, entropy-favored monomers (self folding)^53^. In contrast, our system is governed by homotypic attractions (A binds A, B binds B), where the 1:1 ‘balanced’ composition promotes inter-chain mixing. This analysis reinforces that sequence specific interaction primarily governs polymer associations, while overall sequence composition modulates the extent of mixing but cannot override the effects of patterning.

## IV. DISCUSSION

This study establishes sequence complexity, quantified by bigram Shannon entropy *H*_2_, as a molecular determinant dictating the balance between intermolecular mixing and de-mixing in confined AB-type block copolymers (BCPs). Our systematic study employing exact enumeration of two short BCP chains on a two-dimensional lattice under distinct confinement geometries reveals four critical aspects: i) high-complexity sequences (*H*_2_ = 1.86) not only favor enhanced mixing but also introduce greater heterogeneity in overlapping configurations, particularly under asymmetric confinement (*S*_*r*_). This heterogeneity in *S*_*r*_ arises from distributed short-length repeats, providing entropic flexibility that stabilizes diverse high *N*_*c*_ states, as evidenced by bimodal *D*(*N*_*c*_) distributions [Fig. S2, SI] and multiple conformational clusters in *P* (*R*_end_, *N*_*c*_) (Fig. 5). In contrast, low-complexity blocky sequences (*H*_2_ = 1.26, 1.45) exhibit unimodal, low-mix ensembles, where self-folding via intra-chain contacts yields energetic stability (higher phase-separated contacts) but resists extensive inter-chain overlap [Fig. S1]. ii) Free energy landscapes further reveal this transition as high-*H*_2_ sequences feature surmountable barriers ( 1-2 *k*_*B*_*T*) separating low and high mix minima, enabling cooperative transitions to near-complete overlap. Whereas, low-*H*_2_ landscapes show steep free energy barrier at elevated *N*_*c*_. iii) The high-mixed high complexity sequences undergo co-collapsed state (coupled coil-to-globule transition) where as low *H*_2_ sequences remain expanded and low-mixed across confinements. iv) ‘High *H*_2_ favoring high mixing’ scenario holds also for unbalanced compositions (*A/B* ≠ 1), emphasizing sequence patterning primacy over composition. Fig. 8 illustrates this sequence-dependent behavior: low-complexity sequences (*H*_2_ = 1.26) favor self-folded conformations with minima in low-mixed states, while high-complexity sequences (*H*_2_ = 1.86) undergo cooperative mixing with minima shifted to higher *N*_*c*_.

**FIG. 8.**
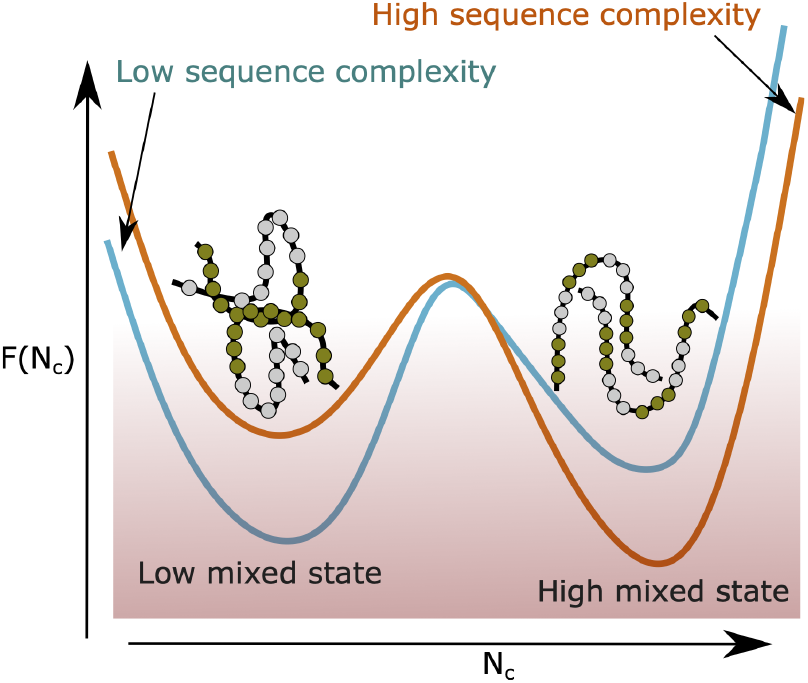
Schematic free energy landscape linking sequence complexity to mixing pathways. Free energy landscape schematic for block copolymers (*H*_2_ = 1.26 and *H*_2_ = 1.86) as a function of inter-chain contacts *N*_*c*_ at *ϵ* = −3. The blue curve illustrates the landscape for *H*_2_ = 1.26 sequence, which exhibit a free energy minimum at low mixing states favoring self folding over mixing due to the blockiness in *H*_2_ = 1.26. The orange curve represents *H*_2_ = 1.86 variants, where the free energy minimum shifts to higher *N*_*c*_ values in the mixed state. The short length repeats in *H*_2_ = 1.86 suppresses intrachain self-folding and promotes intermixing with the neighboring chain.

To assess whether the sequence-complexity regimes explored by our model are relevant to real proteins, we analyzed the bi-gram entropy *H*_2_ of polyXY regions (two-residue low-complexity tracts) across multiple proteomes [Fig. 7]^54^. Remarkably, the distribution of *H*_2_ values is centered around 1.3 *±* 0.3 bits, which overlaps precisely with the range in which our model predicts a transition between self-folded, weakly mixed states and inter-chain mixed, co-collapsed states. This correspondence suggests that many polyXY segments in vivo may be naturally tuned to function near the boundary separating mixed and de-mixed states, enabling cells to regulate condensate assembly and disassembly in response to changing conditions.

Although our enumeration-based framework simplifies the vast complexity of single proteins and condensates on a two dimensional lattice, its strength lies in its exact calculation of the partition function, which provides a precise thermodynamic characterization of the interplay between sequence patterning and confinement. More broadly, the sequence–structure relationship uncovered here reflects a fundamental organizing principle that spans from single-chain folding to higher-order condensate assembly. By employing a minimal but exactly solvable model, we demonstrated that the patterning of interaction sites, not just their overall abundance, is a critical determinant of polymer association.

## 1. Acknowledgement

DM thanks Erik Clarkson for reading the manuscript.

