## Supplementary material for "Sequence Complexity Dictates Polymer Mixing": SI

### Supplementary Information
### Sequence Complexity dictates Polymer Mixing

Dibyajyoti Mohanta[1], Manish Dwivedi[2], Hiranmay Maity[3], Debaprasad Giri[4],

**1** Department of Chemistry, The State University of New York at Buffalo, NY, USA
**2** Department of Physics, Institute of Science, Banaras Hindu University, Varanasi, India
**3** Department of General Science, BITS, Pilani, Dubai Campus, Dubai International Academic City, Dubai, UAE
**4** Department of Physics, Indian Institute of Technology (BHU), Varanasi, India

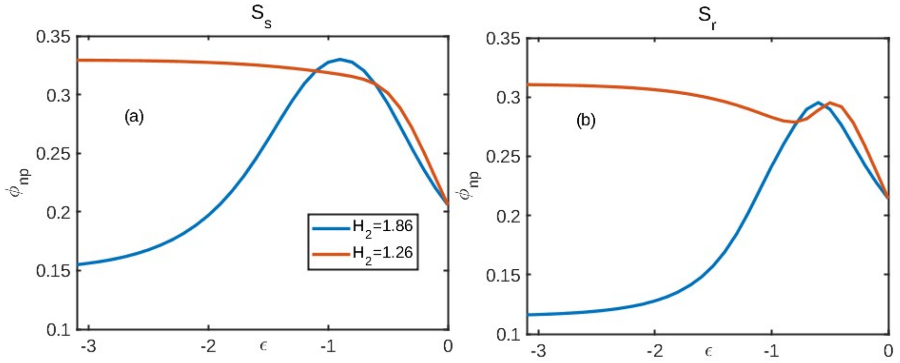


**Fig S1.** Weighted average of fraction of intra contacts per chain $\phi_{np} = N_p/(N+1)$, where $N_p$ is the mean number of intra contacts per chain. Low complexity blocky sequences $H_2 = 1.26$ higher self folded configurations compared to $H_2 = 1.86$ sequences in both confinements.

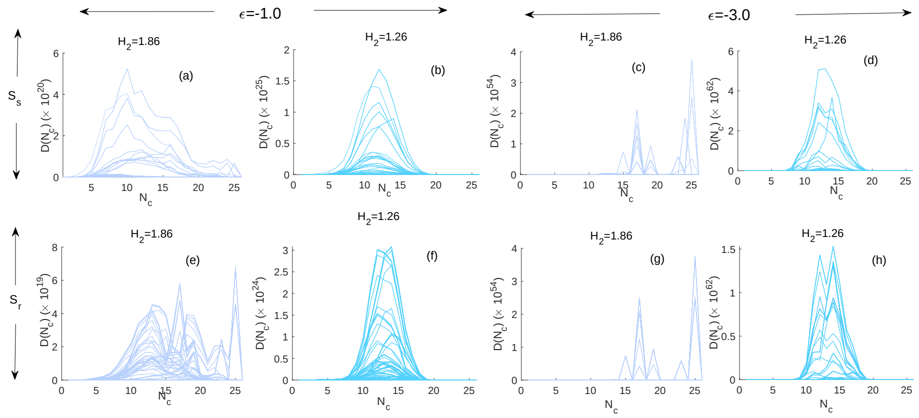


**Fig S2.** Sequence complexity controls thermodynamics of polymer association: Weighted density of states $D(N_c)$ as a function of inter-chain contacts reveals that at $\epsilon = -1.0$ (a,b,e,f), $H_2 = 1.86$ sequences show broad distribution of $N_c$ (a,e) than $H_2 = 1.26$ sequences (b,f). This further accelerates at higher strength $\epsilon = -3.0$ (c,g) for $H_2 = 1.86$, which shows bimodal distributions where as $H_2 = 1.26$ sequences remain unimodal, trapped in less mixed (low $N_c$) states.

The weighted density of states, $D(N_c)$, as a function of inter-chain contacts $N_c\,(= m_{sp} + m_{nsp})$ (Figure S2) reveals fundamental differences in the configurational landscapes accessible to both sequence types.

$$D(N_c) = \sum_{s_{sp}, s_{nsp}} C(m_{sp}, m_{nsp}, s_{sp}, s_{nsp}) \exp(-\beta\epsilon)^{m_{sp}} \exp(-\beta\epsilon)^{s_{sp}} \tag{S.1}$$

At moderate interaction strength ($\epsilon = -1.0$) [Fig. S2 a,b,e,f], both sequence types exhibit relatively symmetric distributions centered around $N_c \approx 15$, indicating thermodynamic equilibrium between low-mixed and partially mixed states within both confinements. However, the distribution for $H_2 = 1.86$ sequences display a broader distribution indicating a wider range of inter-chain configurations, particularly pronounced under $S_r$ where lateral geometric constraints reduce the entropic cost of inter-chain alignment [**?**].

The distinction becomes enhanced at high attraction ($\epsilon = -3.0$), where $H_2 = 1.86$ sequences develop a characteristic bimodal distribution with well-populated peaks corresponding to a dispersed, low-contact state ($N_c \approx 15$) and a highly associated, high-contact state ($N_c \approx 25$) [Fig. S2 c,g]. The broad, bimodal distributions observed for $H_2 = 1.86$ sequences indicate that transitions between dispersed and associated configurations remain feasible without overwhelming barriers [**?**]. In contrast, low-complexity sequences maintain a single unimodal distribution [Fig. S2 d,h] even under strong interactions. The higher amplitude of narrow $D(N_c)$ distribution for $H_2 = 1.26$ sequences reveal the entropic penalty for both fully separated and extensive mixing is large, confining the system to a narrow restricted conformational ensemble where low overlapping dominate [**?**]. These results establish sequence complexity as a molecular switch that controls whether a binary BCP system remains in a low-mixed, self-folded regime or enters a highly overlapped state.

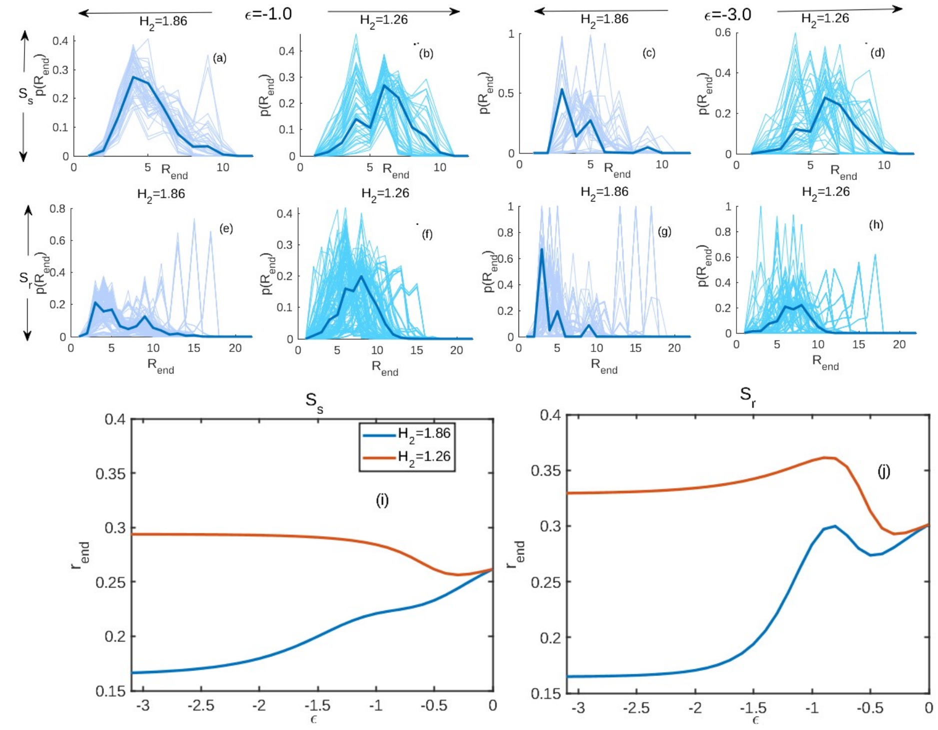


**Fig S3.** Complexity effects size of the polymer: Comparison of end to end distance ($R_{end}$) distribution of polymer 1 between confinements $S_s$ and $S_r$ at interaction strength $\epsilon = -1$ (a,b,e,f) and $\epsilon = -3$ (c,d,g,h) for all origin pairs with the solid line presenting the ensemble average in each case. (i) and (j) represent the average variation of ensemble average end to end distance $r_{end} = R_{end}/(N+1)$ with interaction strength $\epsilon$ Higher sequence complexity $H_2 = 1.86$ system in both confinements attain globule or condensed form while $H_2 = 1.26$ system remain in a kinetically trapped self associated coil structure with higher $r_{end}$.

Sequence dependent differences in intermolecular contacts result in diverse conformational ensembles and size distribution of polymers. To show that, we present the end-to-end distance ($R_{end}$) distribution of polymer 1 in each 2-polymer system for all origin pairs in Fig. S3 (a-h). Under moderate interaction strength ($\epsilon = -1.0$), the distributions begin to differentiate as $H_2 = 1.86$ sequences (Fig. S3 a, e) develop a pronounced shift towards smaller $R_{\text{end}}$ values compared to low-complexity sequences $H_2 = 1.26$ (Fig. S3 b, f) especially. This compression becomes dramatically enhanced at strong interactions ($\epsilon = -3.0$), where high-complexity sequences due to their short repeat sequences undergo multiple simultaneous inter-contacts (multivalency) may favoring cooperative co-collapse of the 2-polymer system (Fig. S3 c, g) . The inter-molecular contact driven co-collapse for polymers in high-complexity sequences is also recently seen on IDP related studies [**?**]. On the other hand, low-complexity sequences maintain coil-like configurations with $\langle r_{\text{end}} \rangle \approx 7-8$ (Fig. S3 d, h) [**?**]. In Fig. S3 (i,j), we concise the size distribution in the form of average scaled end-to-end distance $r_{end} = \frac{R_{end}}{N+1}$ variation with attraction strength $\epsilon$ which shows $H_2 = 1.86$ sequences undergo coil-globule like transition driven by favorable energetics of inter-polymer contact formation in both confinements while $H_2 = 1.26$ sequences do not show any signature of such compaction in size [**?**].
